# Merging in-solution X-ray and neutron scattering data allows fine structural analysis of membrane-protein detergent complexes

**DOI:** 10.1101/324103

**Authors:** Gaëtan Dias Mirandela, Giulia Tamburrino, Miloš T. Ivanović, Felix M. Strnad, Olwyn Byron, Tim Rasmussen, Paul A. Hoskisson, Jochen S. Hub, Ulrich Zachariae, Frank Gabel, Arnaud Javelle

**Affiliations:** Strathclyde Institute of Pharmacy and Biomedical Sciences, University of Strathclyde, Glasgow, G4 0RE, U.K.; Computational Biology, School of Life Sciences, University of Dundee, DD1 5EH, UK; Physics, School of Science and Engineering, University of Dundee, Dundee, DD1 4NH, U.K.; Theoretical Physics, Saarland University Campus E2 6, 66123 Saarbrcken, Germany.; School of Life Sciences, College of Medical, Veterinary and Life Sciences, University of Glasgow, Glasgow, G12 8QQ, U.K.; School of Medical Sciences, University of Aberdeen, Foresterhill, Aberdeen AB25 2ZD, U.K.; Institut Laue-Langevin, 71 Avenue des Martyrs 38042 Grenoble, France; University of Grenoble Alpes, CEA, CNRS, IBS, 38000 Grenoble, France.

**Author notes:** GT and MTI contributed equally to this work. JSH, UZ and FG contributed equally to this work.

**Keywords:** Membrane transporter, Molecular dynamic simulation, Protein biophysical characterization, Small Angle Scattering data analysis, Structural biology, WAXSiS

## Abstract

In-solution small angle X-ray and neutron scattering (SAXS/SANS) have become popular methods to characterize the structure of membrane proteins, solubilized by either detergents or nanodiscs. SANS studies of protein-detergent complexes usually require deuterium-labelled proteins or detergents, which in turn often lead to problems in their expression or purification. Here, we report an approach whose novelty is the combined analysis of SAXS and SANS data from an unlabeled membrane protein complex in solution in two complementary ways. Firstly, an explicit atomic analysis, including both protein and detergent molecules, using the program WAXSiS which has been adapted to predict SANS data. Secondly, the use of MONSA which allows to discriminate between detergent head- and tail-groups in an *ab initio* approach. Our approach is readily applicable to any detergent-solubilized protein and provides more detailed structural information on protein-detergent complexes from unlabeled samples than SAXS or SANS alone.

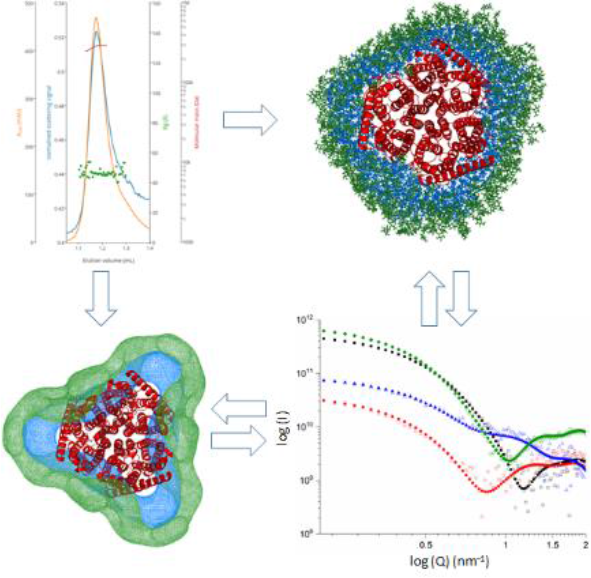

---

Integral membrane proteins form the entry and exit routes for nutrients, metabolic waste and drugs in biological cells, and they are involved in key steps of signaling and energy transduction. They thus play a central role in a variety of biological processes with exceptional medical relevance.^1^ Structural information on membrane proteins has traditionally been obtained by X-ray crystallography aided by detergent molecules that replace the lipids during the purification and crystallization processes. Detergents stabilize membrane proteins by shielding the hydrophobic domains from the aqueous environment.^2^ However, the translocation cycle underpinning membrane transporter activity requires substantial conformational variability and in many cases, the static structural insight achieved by X-ray crystallography has proven insufficient to capture the essential functional information of these systems.^3^ For this reason, there is considerable interest in the application of Small Angle Scattering (SAS) methods to structurally characterize membrane proteins. Recently efforts have been dedicated to developing combined in solution small angle X-ray/neutron scattering (SAXS/SANS) approaches to investigate membrane proteins stabilized by detergent or nanodics.^4–6^ Further developments in these areas have faced important obstacles. Crucially, the electron density of the detergent shell encompassing the hydrophobic domains of membrane proteins differs from the electron density of the protein. Hence, it is difficult to obtain a model of a protein-detergent complex using *ab initio* SAXS-based methods, which typically assumes a uniform electron density across the entire complex. To circumvent this problem, SANS experiments making use of contrast variation either by using deuterium labelled proteins and/or detergent molecules have been employed. However, difficulties are sometimes encountered in the expression and purification of deuterated proteins, as well as the limited availability of deuterated detergents.^4^ To overcome these issues, we report a new methodology that combines SAXS and SANS from unlabeled (i.e. non-deuterated) proteins and/or detergent samples to obtain detailed structural information on protein-detergent complexes. This approach is readily applicable to any detergent-solubilized protein.

We used the ammonium transporter AmtB from *Escherichia coli*, a structurally well-studied member of the ubiquitous and medically important Amt/rhesus family of proteins, to develop and validate our methodology.^7^ To stabilize AmtB, the detergent n-dodecyl-β-D-maltoside (DDM) was used throughout the purification process (Supporting Information). Size exclusion chromatography in-line with multi-angle light scattering (SEC-MALS) analysis showed that the AmtB-detergent complex comprises 285±12 DDM molecules (Figure S1 and Table S1). Independently conducted analytical ultracentrifugation (AUC) experiments revealed a detergent shell of 321±1 DDM molecules (Figure S2 and Table S1). Taken together, these independent findings indicate that the detergent corona around AmtB is likely to include between 260 and 320 DDM molecules.

We next exploited atomistic molecular dynamics simulations of the AmtB-DDM complex and scored the models against SAXS data to resolve the experimental uncertainty regarding the size of the detergent corona. AmtB in the physiologically functional trimeric form (pdb ID: 1U7G)^8^ was simulated surrounded by DDM coronas of 260, 280, 300, 320, 340 and 360 molecules. A representative model obtained for a detergent corona containing 320 molecules of DDM is given Figure 1.

**Figure 1.**
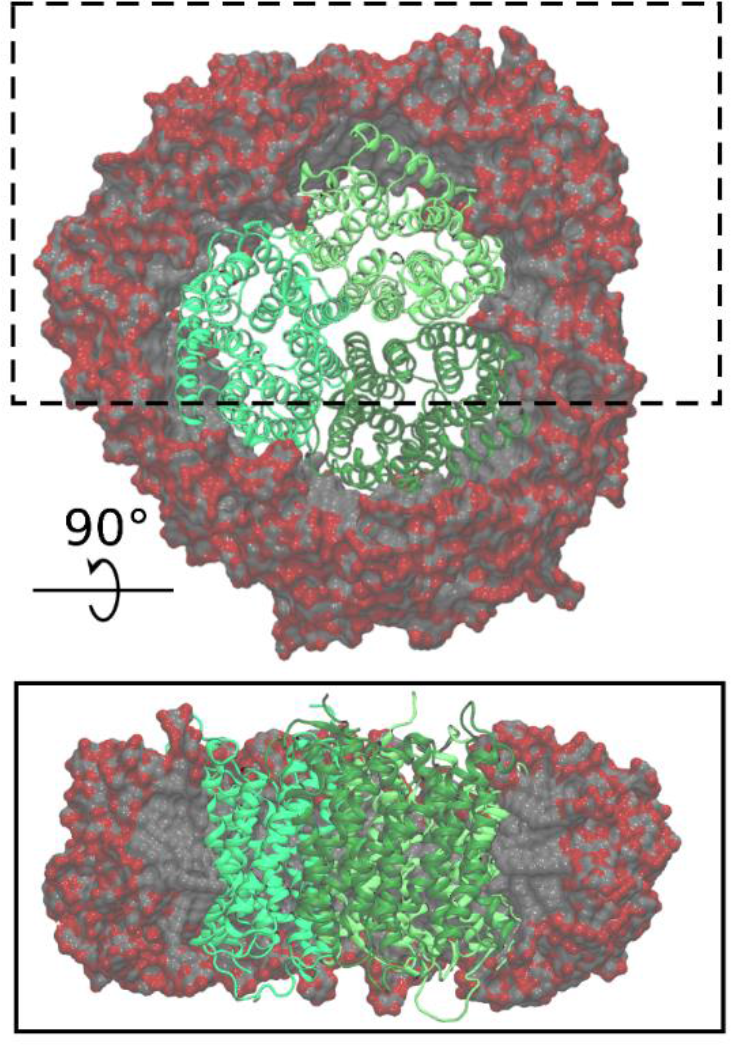
Atomistic model of the AmtB-DDM complex containing 320 DDM molecules. The model displays an equilibrated complex. In the trimer, each AmtB monomer is shown in a different shade of green, and the DDM carbon and oxygen atoms are shown in grey and red, respectively. The upper picture of the panel displays the complex seen from the top, the lower shows a side-view of the complex where the DDM molecules outside of the box highlighted in the top panel are omitted, to illustrate the interior of the micelle.

During the equilibration phase, the DDM molecules adopted the typical toroidal shape reported for other protein-detergent complexes^9–10^, with their hydrophilic heads facing the aqueous solution and their hydrophobic tails oriented toward the inside of the complex (Figure 1). As previously shown, the detergent corona further adapted to the shape of the transmembrane surface of the protein.^10^ Our simulations indicate that the protein-detergent complexes are stable, and although some reorientation of DDM was observed, in particular during the first stages of the simulations, no dissociation of detergent molecules from the protein was detected after 20 ns of simulation time. We next computed SAS curves for the simulated complexes and compared them with experimental SAS measurements (Figure 2–3).

**Figure 2:**
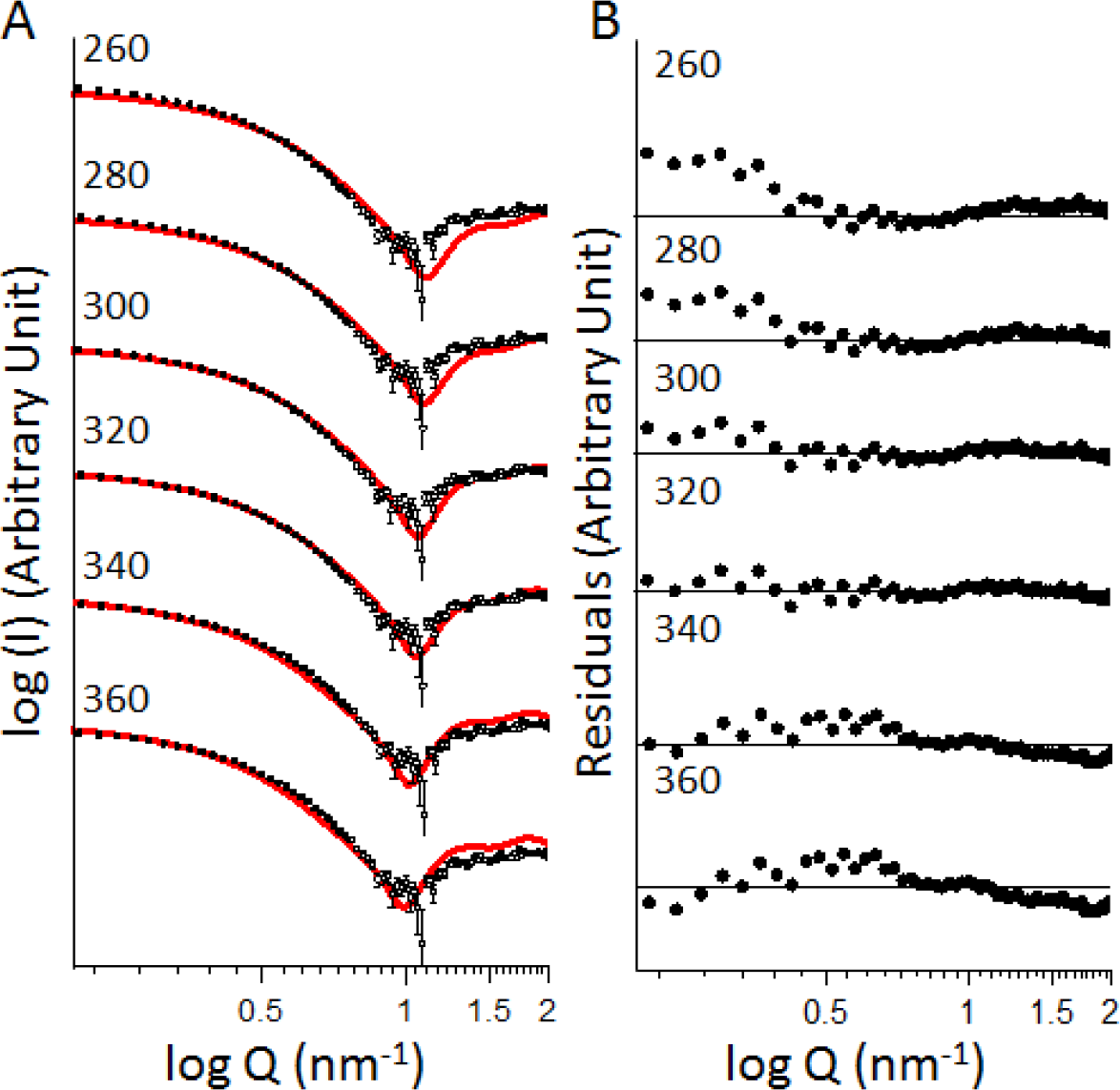
(A) Comparison of the experimental (symbols) and computed (red line) SAXS curves of the AmtB-DDM complex containing between 260 and 360 DDM molecules. For all plots, the maximum and minimum value for Y-axis are 10^11^ and 10^5^. (B) Residual error plot expressed as a the experimental minus computed scattering intensity. For all plots the maximum and minimum value for Y-axis are 40 and −40. *Q* = (4π sin(θ)/λ), where 2θ is the scattering angle

**Figure 3:**
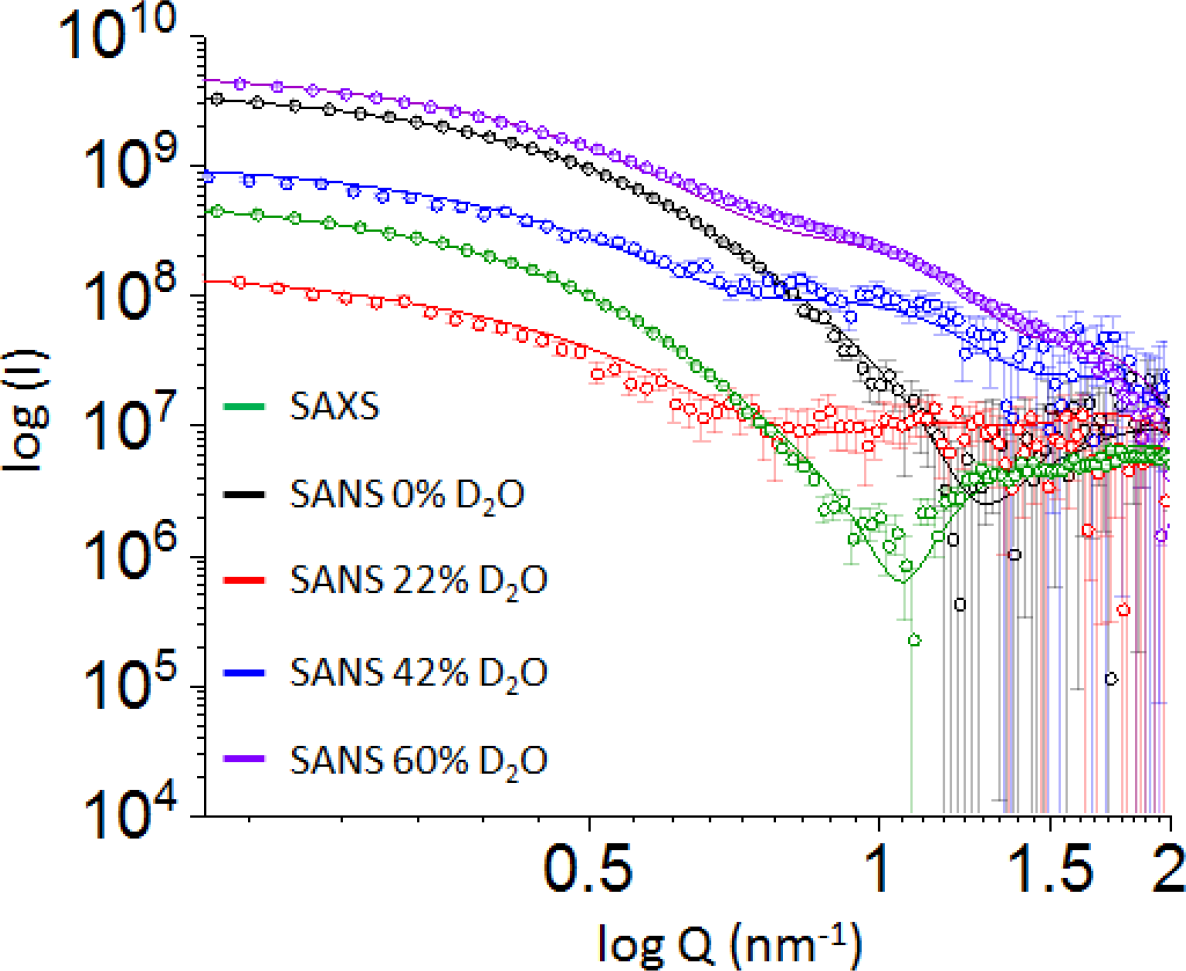
Comparison of the experimental (symbols) and compute (red line) SAXS/SANS curves for the model containing 320 DDM molecules.

It has previously been shown that single structures extracted from MD trajectories do not fully capture the characteristics of the solution ensemble.^9^ We therefore calculated the predicted SAXS curves from conformational ensembles comprising 9000 individual configurations as observed in 70-160 ns simulations of each differently sized complex. The SAXS curves were obtained using explicit-solvent calculations as implemented in the WAXSiS method, thereby taking into account accurate atomic models for both the hydration layer and the excluded solvent, and consequently avoiding any solvent-related fitting parameters (Figure 2).^11–12^

SAS experiments are very demanding in terms of requirements of sample quality^13–14^, therefore, before recording SAS data, we ascertained that our samples was monodisperse and that AmtB was pure, stable and critically, active in detergent (Supporting information, Figure S1-S3). We subsequently collected experimental SAXS data following size-exclusion chromatography of the AmtB-DDM complex. The radius of gyration (Rg) was found to be constant across the elution peak (Figure S1) indicating the monodispersity of the complex and good data quality. Importantly, the scattering curves predicted for the models containing 260, 280, 300, 340 and 360 DDM molecules deviate slightly from the experimental data (Figure 2 and S4). In contrast, the curve computed for the MD model containing 320 DDM molecules was nearly indistinguishable from the experimental SAXS data (Figure 2 and S4). Furthermore, the values for Rg obtained by the Guinier approximation from the experimental data and from for the MD model containing 320 DDM molecules were in quantitative agreement (Table S3). This suggests that the overall dimensions of the protein-detergent complex containing 320 molecules of DDM in simulation and experiment are identical. We therefore identified the AmtB complex with 320 DDM molecules as the predominant species observed in solution. It is important to note that the overall information content of SAXS is relatively low, and thus agreement between experimental and back-calculated curves may be insufficient to serve as unambiguous evidence for a structural model.^15^ Specifically, in the context of a protein-detergent complex, SAXS data reports on the overall shape of the complex, whereas they do not provide independent information on the individual contributions from the protein and the detergent corona. Therefore, we employed SANS together with contrast variation to more firmly validate our computational model.

We collected SANS data at four contrast points (0%, 22%, 42% and 60% deuterated water-D_2_O) to differentiate between the individual components of the protein-detergent complex. To ensure that the samples were stable over the course of the SANS experiment, the hydrodynamic property of the proteins were analyzed before and after the SANS measurements by analytical size exclusion chromatography. No difference were observed in the elution profile, confirming the stability of the protein during the SANS experiment (Figure S5). To ascertain the reproducibility and the quality of our measurements, two independent set of SANS data have been measured using two batches of AmtB purified independently and the two dataset were identical within noise (Figure S6). It has previously been shown that in the absence of D2O in the buffer, neutron scattering from DDM micelles originates primarily from the hydrophilic head groups.^16^ We calculated (Supporting Information) the overall contrast match point of DDM to be at 22% D_2_O, while the contrast match point for typical proteins is around 42% D_2_O.^4, 17^ Consequently, the scattering contribution is dominated by the protein and the DDM hydrophilic head group in a buffer containing 0% D_2_O, by the protein at 22% D_2_O and by the complete detergent corona at 42% D_2_O. To compare the experimental neutron scattering data with the MD-generated models, SANS curves were calculated using WAXSiS for 9000 individual configurations observed during 70-160 ns MD trajectories of each of the complexes. To this end, we extended the WAXSiS method, originally developed for SAXS predictions, to also allow SANS predictions with explicit-solvent models at various D_2_O concentrations (Supporting Information). The experimental curves were fitted to the calculated curves following *I*_fit_ = *f*·*I*_exp_+*c*, thereby accounting for scattering contributions from the incoherent background with the fitting parameter *c*. However, neither the hydration layer nor the excluded volume were adjusted. Congruent with the analysis of the SAXS data, all SANS data sets agreed best with the curves calculated for the model incorporating 320 molecules of DDM (Figure 3 and Figure S7). Hence, the SANS and SAXS data consistently validate our MD model with 320 DDM molecules. Secondly, the excellent agreement we observe between the experimental and calculated SAXS curves shows that the overall organization of the complex is accurately reflected by the atomistic model. Finally, the good agreement between experimental and computed SANS curves indicates that the MD model describes accurately the hydrophobic and hydrophilic phase of the detergent ring as well as the position of AmtB inside the corona.

Importantly, the crystal structure of AmtB was used to produce our MD trajectories, which precludes the possibility of applying this combined MD/SAXS/SANS approach to membrane proteins of unknown structure. We therefore applied, in the final step, an independent “MD-free” approach to obtain a full *ab-initio* model that captures detailed structural information on the complex without using the crystal structure of AmtB. To achieve this, we merged our complete SAXS and SANS data and conducted a multiphase volumetric analysis of the complex using MONSA^18–19^ (Figure 4). Importantly, we introduced two separate phases to describe the head and tail groups of the DDM detergent corona.

**Figure 4.**
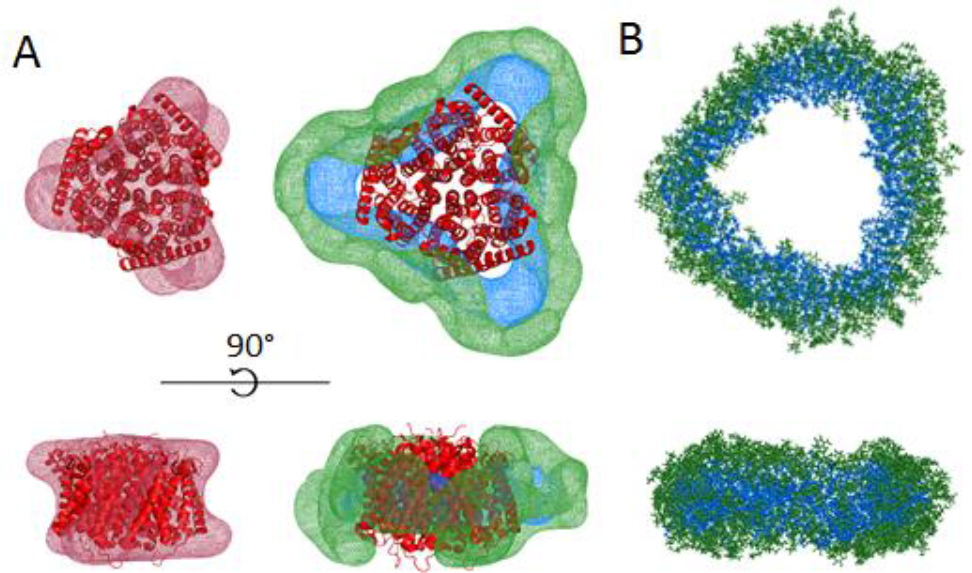
(A) MONSA multiphase modelling using the SAXS and SANS data. (B) Molecular-dynamics generated model of the detergent corona (320 molecules) surrounding AmtB. The phase corresponding to the protein is represented in red mesh, while the hydrophilic and hydrophobic detergent densities are represented in green and blue, respectively.

Assuming the volume of a DDM molecule to be 690 Å^3^ (350 Å^3^ and 340 Å^3^ for the head and the tail respectively)^4^, we imposed a volume of 112,000 Å^3^ and 108,800 Å^3^ for the hydrophilic and hydrophobic phases of the 320 DDM molecules. The volume of AmtB (166,864 Å^3^) was calculated based on its amino acid sequence alone (Supporting Information). Moreover, since the trimeric nature of AmtB in solution was confirmed by our SEC-MALS and AUC data (Figure S1, Figure S2 and Table S1), we also imposed a P3 symmetry of the complex for the analysis. Crucially, all this information can be easily obtained for any membrane protein solubilized in detergent using widely accessible and complementary biophysical techniques (SEC-MALS/AUC in this study). Ten MONSA runs were performed yielding similar *ab-initio* envelopes for AmtB. A representative MONSA model is shown in Figure 4, which faithfully reflects both the size and shape of the MD-generated model. The protein envelope is a good representation of the crystallographic structure of AmtB and is, furthermore, confined inside the detergent corona. Importantly, the joint use of both SAXS and multiple SANS datasets allowed to distinguish the head- and tail-groups of the detergent corona and place them correctly with respect to the protein surface and solvent. Such detailed insight is usually not achieved with *ab initio* models unless additional contact restraints are applied^20^: the detergent ring fits the contours of the protein and the positions of the two detergent phases (head- and tail-groups) are particularly clear. The hydrophobic phase is strictly contained between AmtB and the hydrophilic ring, with only the tails of DDM being in contact with the hydrophobic surface of the transmembrane domain. Hence, without using deuterated protein or detergent, and without information about the 3D structure of AmtB, the combination of SAXS and SANS data capture the essential structural detail contained in membrane-protein detergent complexes in solution.

In summary, there is considerable interest in developing SAS methodology further to allow routine investigation of membrane proteins. We have adapted WAXSiS to account for SANS data and therefore open up this software package for future projects including both types of scattering data. Using our methodology, based upon a combination of SAXS/SANS measurements and MD simulations, we have been able to propose an atomic model of a protein-detergent complex. Our integrative approach demonstrates that combining SAXS, SANS, and iterative simulations provides more detailed structural information than each of the methods alone.

It is widely recognized that cryo-electron microscopy (cryo-EM) will revolutionize the structural analysis of membrane proteins in the near future.^21–22^ It is our believe that an hybrid approach, combining in solution SAS technique, *in silico* modeling and cryo-EM will allow to better track and describe conformational changes of membrane proteins in solution, induced by ligand or cofactor binding. In this context, it was important to account accurately for the bound detergent molecules, which is greatly improved by combining SAXS and SANS data at various contrasts. Secondly, our multiphase analysis, which merges SAXS and SANS data, without using deuterated protein or detergent, allowed us to obtain unprecedented structural information on the phase density of the detergent, in particular to distinguish head- and tail-groups in the assembled membrane protein-detergent complex. This is particularly relevant as deuterated media/detergents are often expensive and toxic for bacteria, leading to decreased protein yields.^23^ Crucially, the multiphase analysis does not require information on the 3D structure of the protein, which opens the possibility to apply this methodology to a wide range of important membrane proteins that have so far remained inaccessible to high resolution structural analysis. Whilst SAS has become a popular technique amongst structural biologists, combinations of SANS, SAXS and MD has remained underexploited by the community. In this context, our work represents a significant advancement in data acquisition, model validation, development of new software and multiphase volumetric analysis to firmly establish SAS technology as a standard method for membrane protein structural biology.

## Supporting Information

Computational and methodological details, as well as three supporting tables seven supporting figures.

## Notes

The authors declare no competing financial interests.

## ACKNOWLEDGMENT

GDM/AJ were supported by a PhD and a Chancellor’s Fellowship from Strathclyde University respectively, GT/UZ acknowledge funding from the Scottish Universities’ Physics Alliance (SUPA). PAH acknowledge the support of the Natural Environment Research Council (NE/M001415/1) and AJ the support of Tenovus Scotland (project S17-07). MTI/FMS/JSH acknowledge support by the Deutsche Forschungsgemeinschaft (HU 1971-1/1, HU 1971-3/1, HU 1971-4/1). We thank the ILL Block Allocation Group (BAG) system for SANS beamtime at D22, Dr A. Martel for help with the setup of the instrument. We thanks Dr. P. Soule (NanoTemper Technologies GmbH) and Dr. M. Tully (DIAMOND, U.K.) for help with the microscale thermophoresis experiments and SEC-SAXS data acquisition respectively.

